# Functional Connectivity Localizes a Distributed Supramodal Core for Naturalistic Viewing

**DOI:** 10.1101/2025.11.01.685026

**Authors:** Viacheslav Fokin, Arefeh Sherafati

## Abstract

Understanding the brain mechanisms that support naturalistic perception is a central goal of cognitive neuroscience. Research has progressed from studying the brain with simple unimodal stimuli to employing richer paradigms that engage visual, auditory, and language modalities in naturalistic environments. Recent multimodal encoding models have shown strong promise in modeling and predicting neural responses to such complex stimuli. Previous studies, however, have primarily focused on two modalities (visual and auditory) and often highlight regions implicated in audiovisual processing (e.g., posterior STS), but do not delineate a compact, replicable subnetwork for multimodal, naturalistic stimuli. While these studies produced valuable maps of sensory integration, their scope was restricted to two modalities and they lacked replication, leaving the full picture of the multimodal integration uncertain. Moreover, no studies have localized a compact, replicable subnetwork via FC-guided ablation under naturalistic viewing. The present study advances our understanding of naturalistic perception by identifying the most informative parcels in Schaefer’s brain atlas that contribute to this task. Building on our earlier work demonstrating that functional connectivity can reveal parcels critical for naturalistic perception, we refine this approach by applying “top drop/bottom drop” and “top keep/bottom keep” selection strategies as well as “broken-stick” analysis to focus on regions most relevant for multimodal integration during naturalistic viewing.

## Introduction

Understanding naturalistic perception and how the brain integrates rich, real-world sensory input has long been a central goal of cognitive neuroscience. Early research relied on controlled experimental paradigms with simplified stimuli, but these approaches captured only limited aspects of naturalistic processing. More recently, deep neural networks have emerged as powerful tools for modeling brain responses to naturalistic stimuli. However, because multimodal integration occurs early in perception and shapes subsequent processing (Ursino, Cuppini, & Magosso, 2014; Hu & Mohsenzadeh, 2025), accounting for it is crucial to fully understand naturalistic perception.

Initial neural network models were unimodal, predicting brain responses to either auditory or visual input alone. These unimodal models proved insufficient, as they could not capture the inherently multimodal nature of naturalistic perception. Studies introducing multimodal models demonstrated that incorporating additional modalities consistently improved predictive performance, with models using visual, auditory, and language inputs outperforming unimodal and bimodal approaches (Fu, Chen, Zhang, Zhang, & Wang, 2025; Oota et al., 2025). Oota and colleagues further showed that across most cortical regions, multimodal models predicted brain activity more accurately than unimodal ones, except in the lateral occipital complex (LOC), a region specialized for object-visual processing. Another study showed that deep multimodal models can successfully predict brain responses to naturalistic audiovisual stimuli (Khoshla et al., 2021).

Prior work has demonstrated that naturalistic brain responses are highly predictable from audiovisual and multimodal inputs using deep neural networks (Khosla et al., 2021; Fu, Chen, Zhang, Zhang, & Wang, 2025) and highlight the role of frontal and higher-order cortical regions in audiovisual integration (Zhou et al., 2025). Yet, most multimodal encoding studies treat the brain as a uniform prediction target, leaving unresolved which specific subsets of parcels are essential for accurate prediction and whether a compact, stable subnetwork could explain most of the variance.

In a previous work (Fokin, V & Sherafati A, 2025), we applied a conservative but functionally meaningful mask that retained only parcels with functional connectivity (FC) above the global median from the Schaefer’s brain atlas. Models trained on this masked subset outperformed those trained on the whole brain, supporting the hypothesis that average FC can serve as a proxy for a parcel’s relevance to naturalistic viewing.

Functional connectivity refers to coordinated activity of brain regions that may be spatially distant but functionally linked (Van Den Heuvel & Pol, 2010). Not all parcels participate in networks engaged by naturalistic movie stimuli; regions unrelated to the task may therefore dilute predictive power. Previous work has used FC to predict cognitive traits and clinical outcomes when combined with deep neural networks (Abrol, Fu, Du, & Calhoun, 2019; Van Wingen, 2025). In addition, the naturalistic viewing task has proved to be an alternative way for mapping the FC networks, thus underscoring the importance of using FC networks and their interplay in understanding naturalistic perception (Vanderwal et al., 2017; Gal, Coldham, Tik, Bernstein-Eliav, & Tavor, 2022), reinforcing its utility for studying multimodal integration.

Building on this foundation, the present study examines the relationship between the absolute value of FC and a parcel’s predictive R-score. We propose a novel method to identify parcels from the Schaefer’s brain atlas, most critical for naturalistic perception. We do so, by analyzing the FC-to-R-score curve to locate a tipping point that separates the most informative parcels from less relevant ones. We use this approach to define a core subnetwork of inter-correlated parcels that may constitute a putative multimodal integration zone. The resulting network is evaluated against known regions implicated in multimodal integration and naturalistic viewing. The present work (i) refines our understanding of how core FC networks drive naturalistic perception, (ii) uncovers previously unrecognized regions contributing to multimodal processing, and (iii) underscores the functional roles of previously known regions in naturalistic perception.

## Methods

### Data and task

We used data from the Algonauts 2025 Challenge, which is derived from the CNeuroMod project—an extensive single-subject fMRI dataset collected during both controlled and naturalistic tasks (Gifford et al., 2025; Boyle et al., 2023). The relevant subset includes two naturalistic movie-viewing datasets: (i) the *Friends* dataset, in which participants watched all episodes from the first six seasons of *Friends* during fMRI scanning, and (ii) the CNeuroMod Movie10 dataset, where participants viewed three feature films and one documentary.

The stimuli consisted of movie visual frames, audio tracks, and time-stamped language transcripts. The neural data included whole-brain fMRI responses from four participants (sub-01, sub-02, sub-03, and sub-05), normalized to the Montreal Neurological Institute (MNI) spatial template (Brett et al., 2002). For analysis, voxel-wise fMRI signals were aggregated into 1000 functionally defined parcels using the Schaefer atlas (Schaefer et al., 2018) resulting in a single time series per parcel.

### Preprocessing

We adopted the preprocessing pipeline provided by the Algonauts 2025 competition.

The whole-brain BOLD fMRI responses were preprocessed using fMRIprep 20.2.x (Esteban et al., 2018) and projected to the Montreal Neurological Institute (MNI152NLin2009cAsym) standard space (Brett, 2002). Functional images were then parcellated by averaging voxel-wise BOLD signals within each of the 1,000 parcels of the Schaefer atlas (Schaefer et al., 2018), resulting in a single fMRI time series for each parcel. Each parcel’s activation was z-scored across sessions (each approximately 15 minutes in duration).

### fMRI time-series and feature alignment

Parcel time series were z-scored over time within each movie block. To account for hemodynamic lag, all stimulus features were shifted with a causal HRF lag (TR = 1.49 sec) of 3 TRs (4.47 sec) (robustness checked for 1–5 TR), and trimmed by 5 TRs at block boundaries to minimize edge effects. The TR value was chosen by training a model with different TR values and choosing the value with which the model performed the best (Table 1). The dataset was trained on Friends seasons 1–5 and Movie10 dataset and tested on Friends season 6. This split was kept identical across all models.

**Table 1:** TR selection testing results.

| HRF delay | Mean R |
| --- | --- |
| 1 | 0.189060 |
| 2 | 0.195425 |
| 3 | 0.196626 |
| 4 | 0.177665 |
| 5 | 0.143377 |

### Functional connectivity and absolute FC parcel score

A subject-level 1000×1000 FC matrix was computed on training data using Pearson correlation between all pairs of parcel time series. Parcel “strength” (or absolute FC score) is defined as the sum of absolute FC weights:

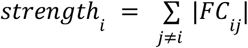

Parcels were ranked by strength, descending for “top” sets and ascending for “bottom” sets.

### Absolute connectivity use robustness evaluation

To assess whether weak correlations could bias our analysis, we excluded low correlation values from the correlation matrix. Because p-values are unreliable in this context, given the large number of data points and resulting inflation, we instead applied several thresholds based on correlation strength. We then evaluated whether the ablation curves changed substantially across thresholds, which would indicate that the results of the analysis were sensitive to weak correlations. We created curves for the top 80%, 90%, and 95% of correlations (p80, p90, and p95) and saw that the function behaviour did not change meaningfully (Figure 1). The shape and behavior of these curves remained largely consistent across thresholds, indicating that the overall trends are robust. Therefore, we can safely work with the full data (no zeroing of connections) and treat thresholding as a robustness check rather than a required preprocessing step.

**Figure 1:**
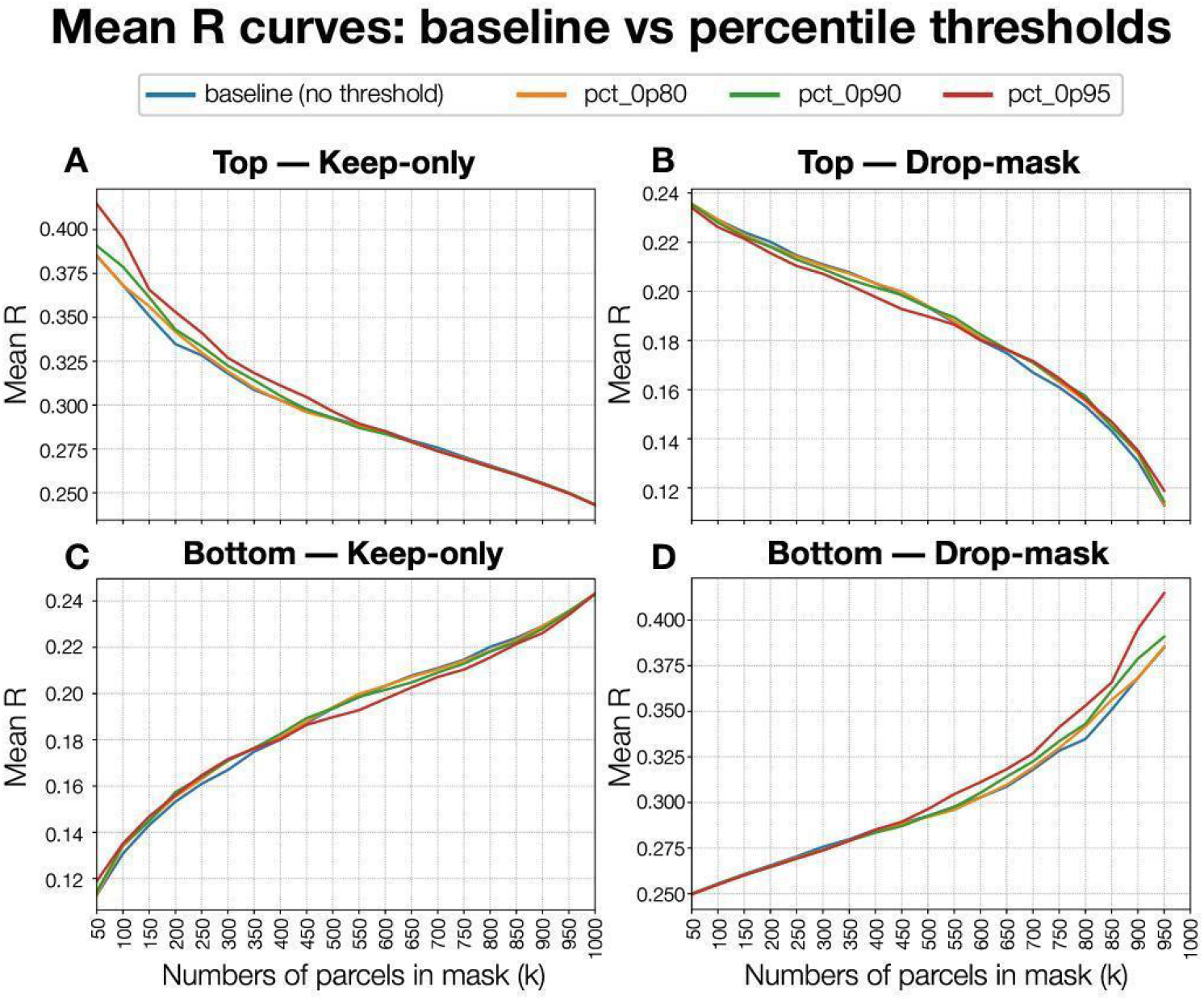
Mean R curves across percentile thresholds. Comparison of mean correlation (R) curves under different correlation strength thresholds to assess robustness of ablation analysis. Each panel shows the mean prediction accuracy (Mean R) as a function of the number of parcels retained (k). for both *keep-only* **(A, C)** and drop-mask **(B, D)** strategies applied to top **(A, B)** and bottom **(C, D)** parcel subsets. Colored lines represent thresholds keeping only the top 20%, 10%, or 5% of correlation values (p80, p90, p95) compared to the unthresholded baseline. Across all conditions, the overall shape and behavior of the curves remain consistent, indicating that removing weak correlations does not meaningfully alter model performance and validating that full (unthresholded) data can be used reliably.

### Encoding and predictive modeling

We used a banded ridge regression model with L2 regularization (Dupre la Tour et al., 2022; Small et al., 2024), trained on 250 principal components (PCs) of multimodal stimulus features extracted from pre-trained visual, audio, and language models. We standardized all stimulus predictors using StandardScaler (z-scoring each feature to zero mean and unit variance). The ridge regularization strength α was implemented via the Adam optimizer’s weight decay parameter (`weight_decay`), which is mathematically equivalent to L2 (Ridge) regularization on the model weights. To respect large-scale functional organization, the 1000 cortical parcels were partitioned into 17 atlas-defined networks (Schaefer 17-network parcellation), and the model employed band-specific readouts aligned with these 17 groups. The visual features were extracted from slow_r50, a 3D ResNet (He et al., n.d.) pretrained on Kinetics-400. We used activations from the *blocks*.*5*.*pool* layer, which captures high-level spatiotemporal video patterns. Language features were obtained from movie transcripts, embedded using BERT (bert-base-uncased) (Devlin et al., 2018). Audio features were derived from the movie’s audio tracks as Mel-frequency cepstral coefficients (MFCCs) (Davis & Mermelstein, 1980). All feature frames were segmented into 1.49-second non-overlapping chunks to match the fMRI repetition time (TR).

Models were trained on the training split and evaluated on the held-out validation split (See Data and Tasks section for details). Model performance was quantified by the mean parcel-wise Pearson correlation *r*_*i*_ between predicted and observed responses on the validation set. Figure 2 presents the overall experimental workflow.

**Figure 2:**
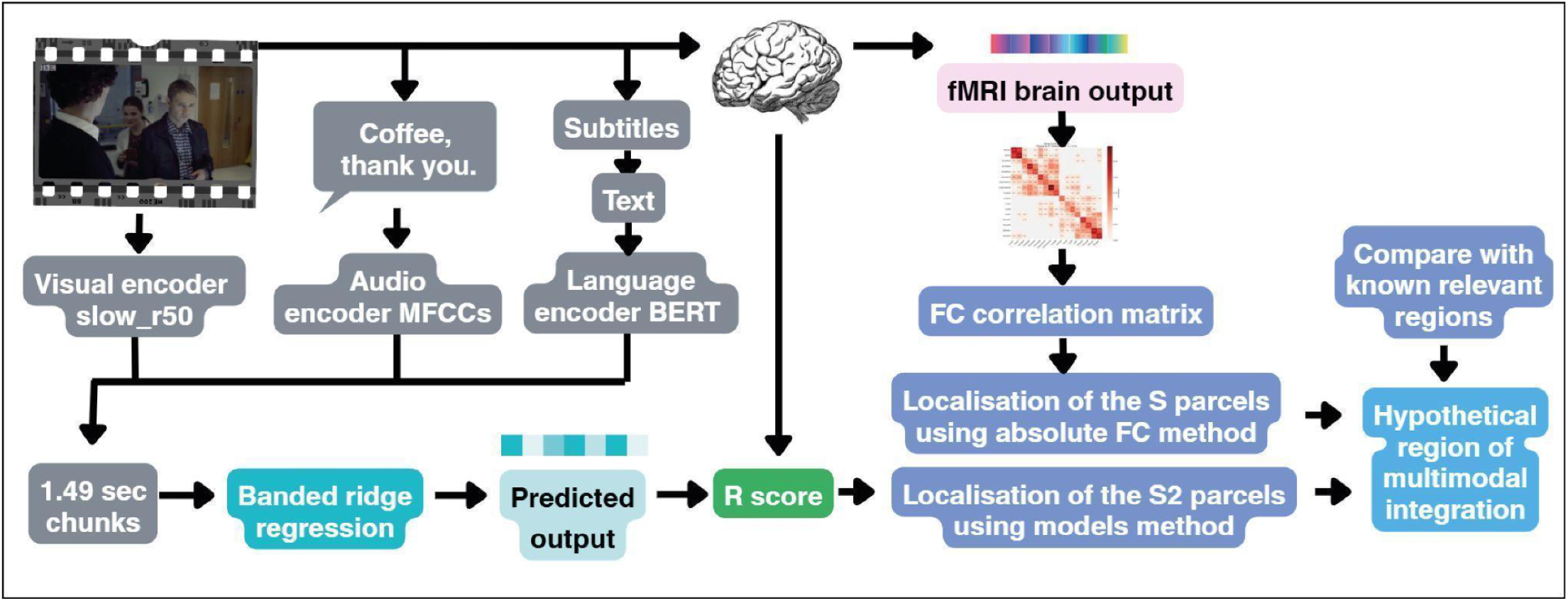
Experiment scheme.

### Piecewise broken-stick analysis

To identify the threshold at which parcels most critical to naturalistic perception begin to dominate predictive performance, we conducted a piecewise (“broken-stick”) regression analysis on ablation curves *R*(*K*), where k indexes parcels ranked by absolute FC.

These curves typically show an initial phase of rapid performance gain followed by diminishing returns. We modeled this using a two-segment linear function:

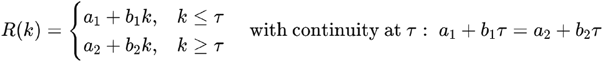

A grid search over candidate breakpoints *τ* (bin size = 50 parcels) was performed. For each *τ*, we fitted two lines using ordinary least squares (OLS) and computed the sum of squared errors *SSE*(*τ*). The optimal breakpoint was chosen as the *τ* minimizing *SSE*(*τ*).

The minimal sufficient set S_FC_ was defined as the smallest *K* at which *k* is closest to the optimal breakpoint.

## Results

### Model Performance with Functional Connectivity Masking

In a previous work (Fokin, V & Sherafati A, 2025), we presented a multimodal encoding model integrating visual, auditory, and linguistic features within the brain’s intrinsic functional architecture to predict neural responses during naturalistic movie viewing. FC correlations were split by an over the median mask (Figure 3A). By constraining feature-based modeling to a functionally defined cortical mask, we achieved a 45.4% improvement in prediction performance, providing strong evidence for the hypothesis that average FC strength reflects a parcel’s relevance to the naturalistic viewing task (Figure 3B).

**Figure 3:**
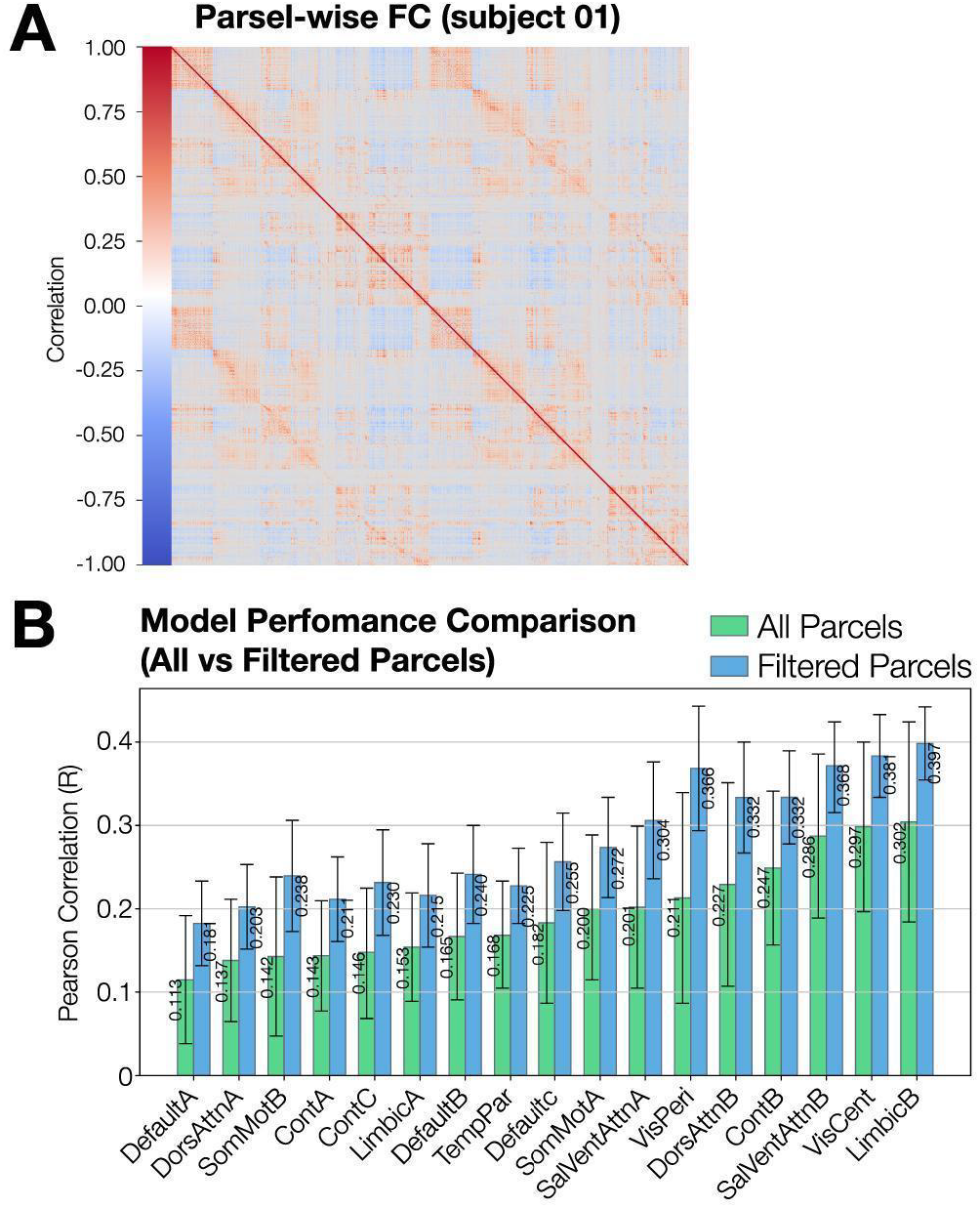
Filtering by FC-derived core parcels boosts model accuracy across cortical networks. **(A)** Parcel-wise functional connectivity (FC) matrix for one representative subject (sub-01), showing correlation patterns among 1000 Schaefer parcels. Warm colors indicate positive correlations and cold colors negative ones, revealing clear modular organization and cross-network coupling across cortical systems. **(B)** Comparison of model prediction accuracy (mean Pearson *R*) between all parcels (green) and the filtered subset (blue) across 17 functional networks. Bars show the mean R per network, and error bars represent the spread of R score within each network. Filtering to the subset parcels consistently increases model performance-especially in visual, control, and salience networks-demonstrati ng that FC-defmed core regions capture the most informative cortical nodes for multimodal fMRI prediction. (Fokin, V & Sherafati A. 2025).

Performance peaked in expected regions, including early visual cortex, bilateral STS, and language-sensitive frontal areas, highlighting model sensitivity to both low- and high-level representations (Fokin, V & Sherafati A, 2025).

### Parcel strength-based ablation analysis

We evaluated the modality-specific contributions across networks to investigate whether improvements were driven by FC-informed selection rather than random parcel removal. We did so by conducting a strength-based ablation analysis.

For each subject, fMRI parcel-wise data were processed as described in *Methods*, and a group-level FC matrix was computed. Each parcel’s absolute FC strength was calculated as the sum of its absolute Pearson correlations with all other parcels. Parcels were then ranked by FC strength.

We defined four masking regimes at increments of *k* ∈ {50, 100, …, 1000}:

- **Top–Keep:** retain the *k* strongest parcels (Figure 4A)
- **Top–Drop:** remove the *k* strongest parcels (Figure 4B)
- **Bottom–Keep:** retain the *k* weakest parcels (Figure 4C)
- **Bottom–Drop:** remove the *k* weakest parcels (Figure 4D)

**Figure 4:**
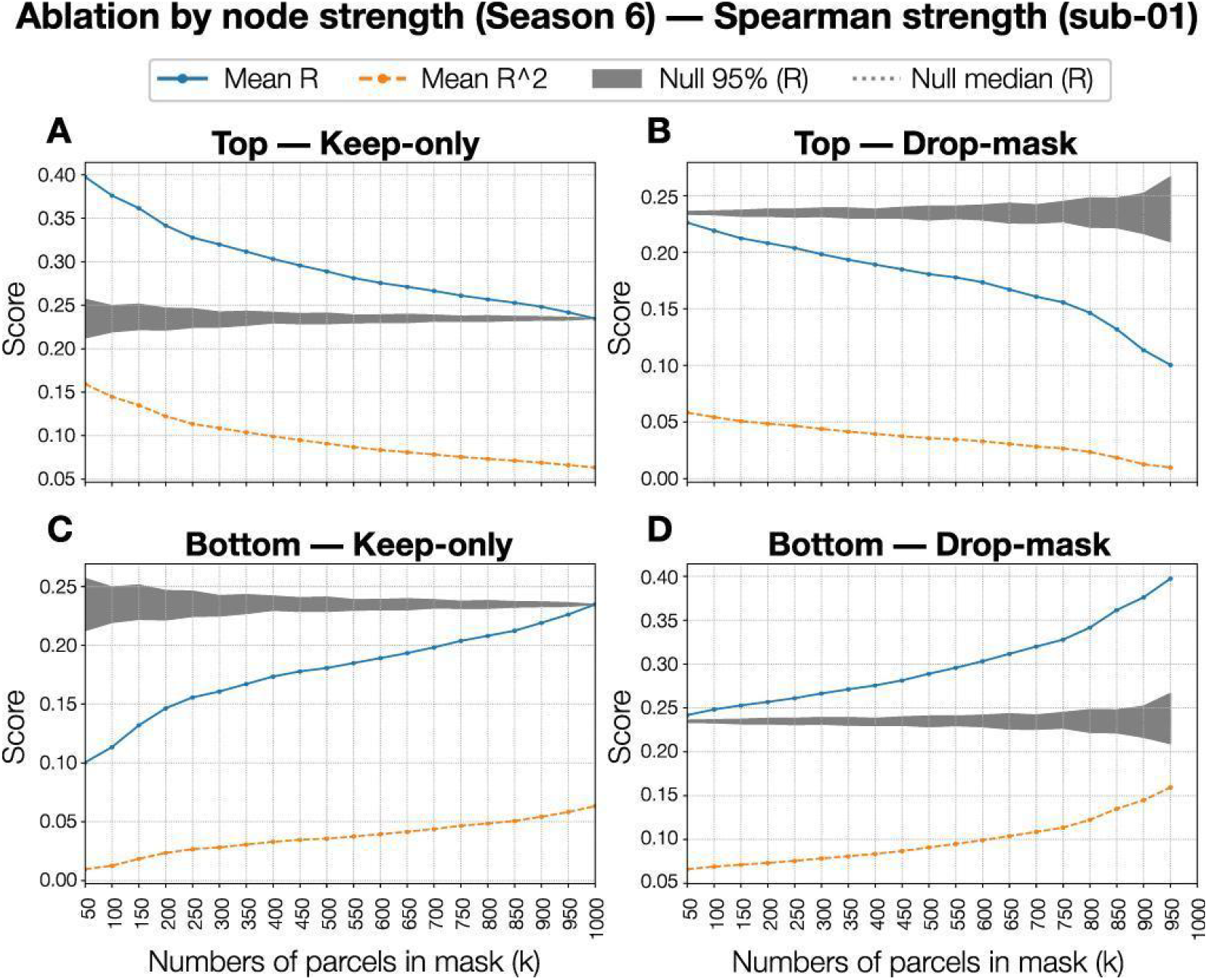
Ablation by node strength reveals the impact of functional hubs on model performance. Plots show the effect of selectively masking parcels based on their functional connectivity (FC) node strength in one representative subject (sub-01, Season 6) Each curve reports the mean Pearson correlation *R* (blue) and mean *R*^2^ (orange) across parcels as a function of mask size (*k*). Gray bands denote the 95% null range and dashed lines the null median. Ablating the strongest parcels (top-drop) causes a marked decrease in model accuracy, whereas retaining only those parcels (top-keep) preserves most predictive power. In contrast, removing the weakest parcels (bottom-drop) slightly improves performance, while retaining only them (bottom-keep) results in substantial loss of predictive capacity. These complementary trends indicate that high-strength FC hubs contnbute disproportionately to stimulus-evoked fMRI prediction, while weakly connected regions play a limited or redundant role.

For each regime, a mean prediction score *R*(*k*) was computed over the remaining parcels (Figure 4).

As a baseline, we generated 100 random parcel masks of size *k* to estimate null distributions (gray lines in Figure 4). Sharp performance declines in Top–Drop and gains in Bottom–Drop confirmed that FC-informed selection captures meaningful task-relevant signals. The null hypothesis—that improvements stemmed merely from removing parcels—was then rejected.

The curves revealed non-linear contributions across parcels: a subset of strongly task-relevant parcels contrasted with parcels that were weakly or negatively predictive. The pronounced change in slope at lower *k* suggested the existence of a core subnetwork driving naturalistic perception.

### Piecewise “Broken-Stick” Analysis

To identify the threshold of core parcel importance, we applied piecewise (“broken-stick”) regression to the Top–Keep curve *R*(*k*) (See *Methods*).

R(k) was smoothed (Savitzky–Golay for keep-curves; isotonic regression for bottom-drop) to reduce staircase artifacts and noise before breakpoint fitting (Figure 5C). The optimal elbow consistently emerged between k ≈ 230–250. We adopted the median estimate (≈240) and rounded to k = 250 to align with the 50-parcel step size, yielding a stable and reproducible threshold for defining the core set S_FC_., visualised on the cortical surface in figure 5A. The same algorithm was also run for top-drop (Figure 5D), bottom-keep (Figure 5E), bottom-drop (Figure 5F).

### Retraining on Core Parcels

To assess whether restricting models to S_FC_ improved predictive power, we compared:

**Figure 5:**
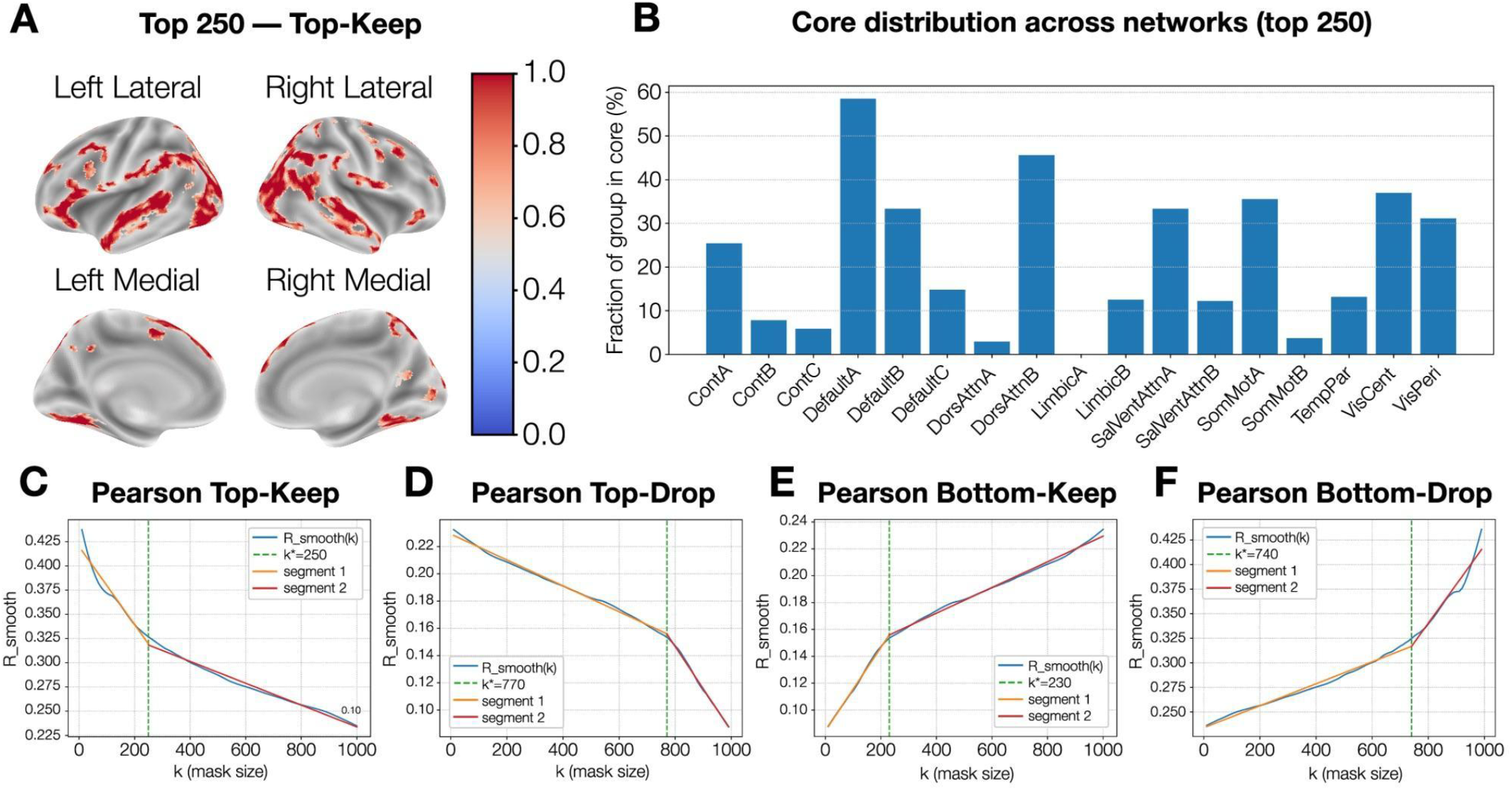
Characterization of the S core within the top 250 functionally connected parcels. **(A)** Spatial distribution of the top 250 parcels identified by the top-keep analysis, projected on cortical surfaces. Color intensity (red-blue scale) reflects the cross-model consistency of each parcel’s inclusion in the S_FC_ core. The core regions concentrate in lateral prefrontal, temporal, and posterior parietal cortices—canonical hubs of multimodal integration. **(B)** Distribution of S"-core parcels across Schaefer-400 functional netv/orks, expressed as the fraction of each network represented in the core. The Default-A, Dorsal-Attention-B, and Visual-Peripheral networks dominate the composition, suggesting that high-strength functional hubs bridge perceptual and control systems. C. Smoothed Pearson correlation (*R* □ □_00_ □ □ (*k*)) curves from the top- and bottom-mask analyses (keep and drop variants). Dashed vertical lines mark the segmentation-based tipping points (k_★_ ≈ 250, 770, 230, 740), separating high-impact and low-impact parcels, These inflection points define the empirical boundaries of the S_Fc_ core and its periphery. **Functional network abbreviations**. VisCent - Visual Central Network; VisPeri - Visual Peripheral Network; SomMotA - Somatomotor A Network; SomMotB - Somatomotor B Network; DorsAttnA - Dorsal Attention A Network; DorsAttnB - Dorsal Attention B Network; SalVentAttnA - Salience/Ventral Attention A Network; SalVentAttnB - Salience/Ventral Attention B Network; LimbicA - Limbic A Network; UmbicB - Limbic B Network; ContA - Control A Network (Frontoparietal); ContB - Control B Network (Frontoparietal); ContC - Control C Network (Cingulo-parietal); DefaultA - Default Mode A Network; Defaults - Default Mode B Network; DefauttC - Default Mode C Network; TempPar - Temporal-Parietal Network.

1. A model trained on all parcels but evaluated only on S_FC_
2. A model retrained exclusively on S_FC_

The S_FC_-trained model showed a significant improvement of ΔR = 0.049 (∼16.5%) in mean parcel-wise correlation compared to the all-parcel model, demonstrating that focusing on core parcels enhances predictive accuracy.

### Modality-specific models and supramodal cores

Our analysis has supported the hypothesis that the average connectivity score enables finding the most relevant parcels for the naturalistic viewing task. Here, we use the average connectivity to gain more insights into the brain networks underlying modality integration and naturalistic viewing.

Matched encoders were trained separately for audio (a), visual (v), language (l), all bimodal pairs (av, al, vl), and the full trimodal model (all). For each model, parcels were ranked by predictive r, ablation curves R(k) were constructed, and piecewise breakpoints determined modality-specific cores S_mod_. For each model, parcels were ranked by that model’s predictive r_j_, constructed R(k) by accumulating the top-ranked parcels in bins of 50, and applied the same piecewise linear regression, single breakpoint method. For each model, we found a tipping point and defined a subset of the most important parcels as S_mod_. The supramodal core was defined as the set intersection across all seven S_mod_ and the S_FC_, visualised on the cortical surface (Figure 6B).

**Figure 6:**
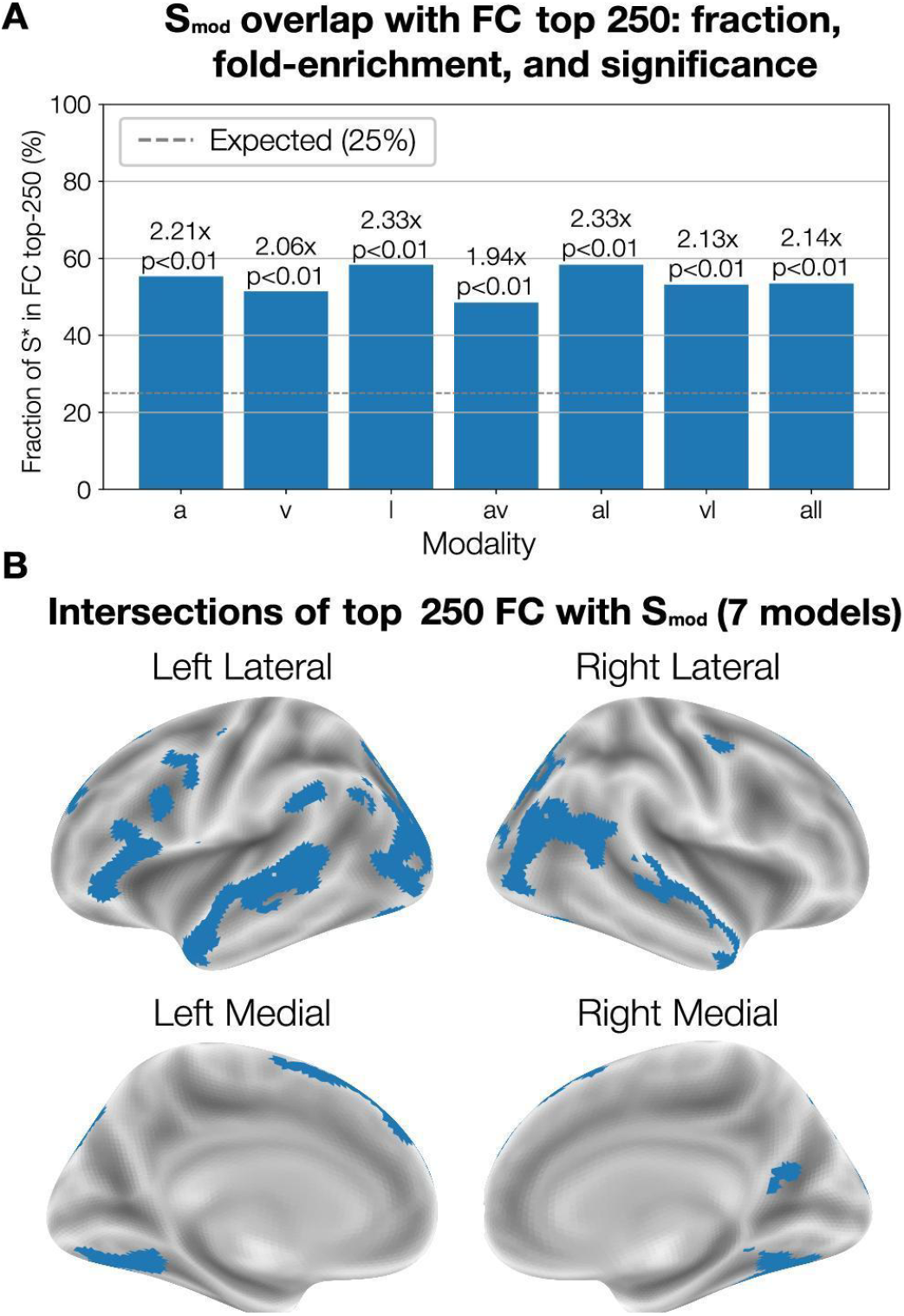
The supramodal core concentrates in the strongest-FC parcels. **(A)** Fraction of parcels in the supramodal set S_sup_, that fall within the Top-250 FC parcels for each model family (a, auditory; v, visual; I, language; av, audiovisual; al. audio-language; vl, visual-language; all, full multimodal). Bars are annotated with fold-enrichment over chance and P-values; the dashed line marks the random expectation (25%). All model families show significant enrichment (*P* < 0.01), with the multimodal models exhibiting the largest effects (≈2.1−2.3×). (B) Lateral and medial cortical views of the supramodal core, defined as the intersection of S_sup_ across seven models with the Top-250 FC set. revealing bilateral clusters in the association cortex.

Fractional overlap and fold enrichment were computed as:

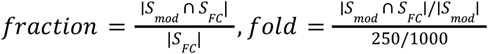

with significance tested using a one-sided hypergeometric test and FDR correction. Overlaps exceeded random expectation (dashed line at 0.25; Figure 6A).

The strongest overlaps occur for language-based and multimodal models (l, al, all), indicating that S_mod_ cores preferentially coincide with highly connected parcels that integrate information across sensory and linguistic systems.

### Comparison to Classical Audiovisual Integration Regions

The supramodal core showed substantial overlap with known multimodal integration regions—notably pSTS, TPO, pSTG, LOC, and MTG—confirming the validity of our FC-based approach. Specifically, comparison to the classical pSTS audiovisual hub revealed a 50% overlap (5/10 parcels) (Figure 7), suggesting that pSTS alone does not fully account for multimodal integration, consistent with prior evidence (Hocking & Price, 2008).

**Figure 7:**
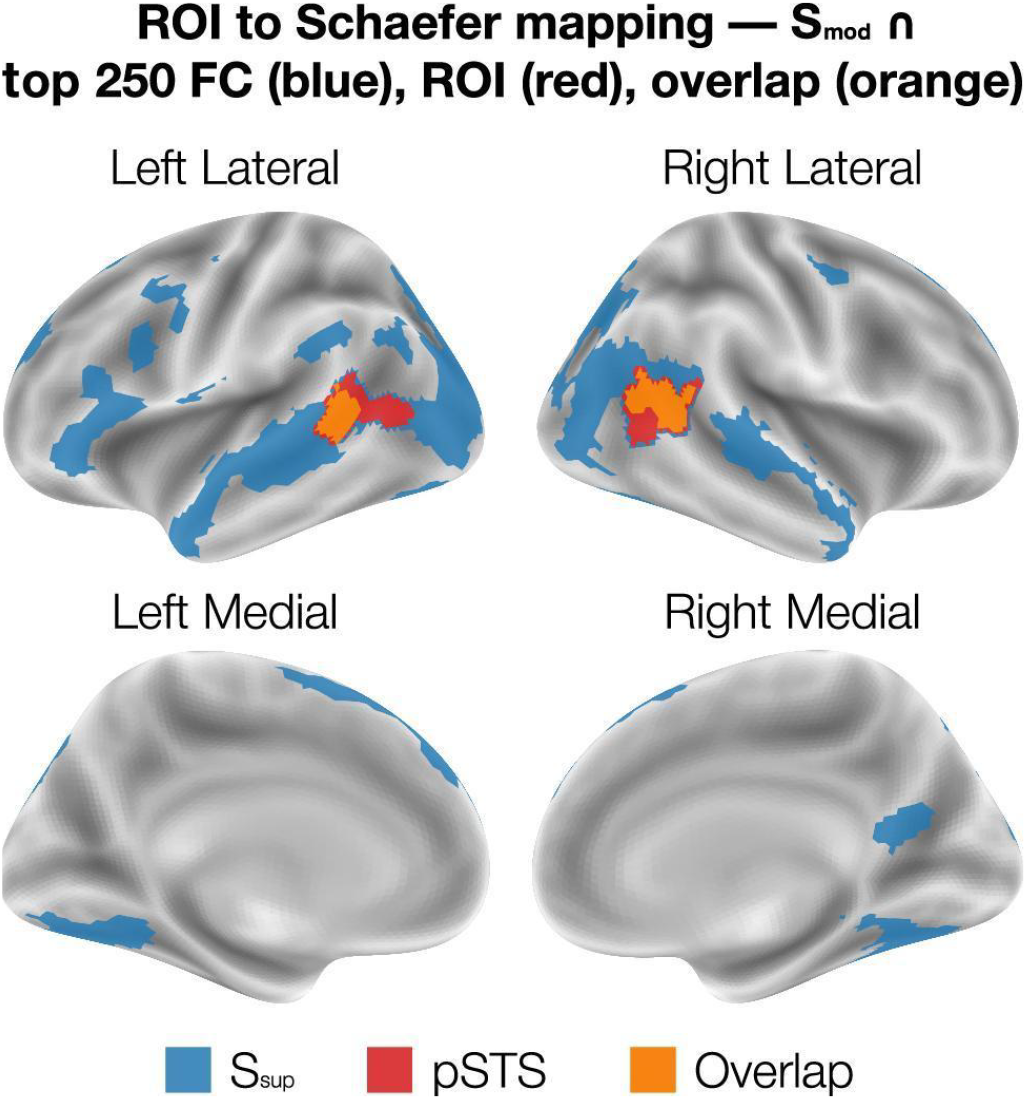
Partial convergence of the supramodal core with the classical pSTS audiovisual ROI. Lateral and medial cortical views show the supramodal core (blue; S_sup_ from the full multimodal model, mapped to the Schaefer atlas), the literature-defined pSTS audiovisual ROI (red), and their overlap (orange). The intersection indicates that pSTS is indeed recruited during naturalistic perception, consistent with prior reports, yet the supramodal core extends well beyond pSTS across association cortex-supporting a distributed, rather than pSTS-exclusive, substrate for multimodal integration.

In order to accurately compare the pSTS region to S_sup_ we mapped ROI coordinates (in MNI space) to Schaefer-1000 (7 networks) parcels using nearest centroids from subject-specific segmentations. For each parcel, we computed the centroid as the mean MNI coordinates of all voxels with that label. Each ROI coordinate was assigned its top-5 nearest parcels by Euclidean distance. We then quantified the overlap between the ROI-derived parcel set and the S_sup_ set using a 2×2 contingency table (Table 2) and exact tests. Specifically, we computed Fisher’s exact test (two-sided) and a hypergeometric right-tail p-value for the observed intersection size, and validated significance with a permutation test (10,000 iterations) that preserved set sizes (|pSTS| and |S_mod_ ∩ S_FC_|). We also report effect-size metrics (Jaccard, precision, recall).

**Table 2:**
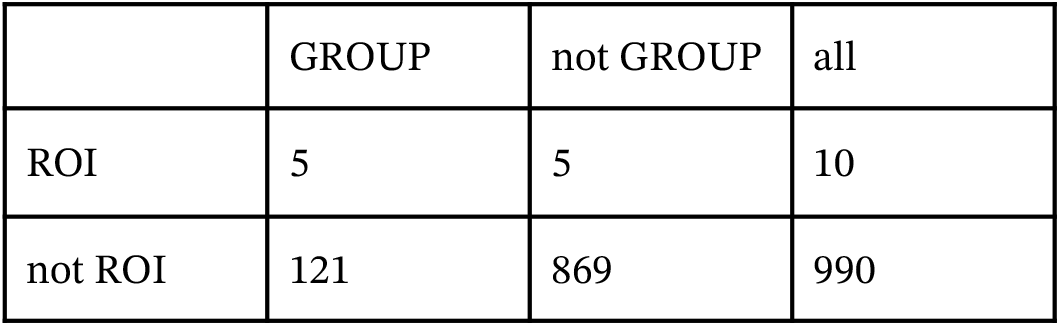
ROI-Group overlap. Set sizes: |ROI| = 10, |GROUP| = 126, Observed overlap = 5; Effect sizes; Jaccard = 0.0382, Precision = 0.0397, Recall = 0.50; P-values: Fisher (two-stded) = 4.38e-03; Hypergeom (right-tail) = 4.38e-03; Permutation = 4.40c-03.

|  | GROUP | not GROUP | all |
| --- | --- | --- | --- |
| ROI | 5 | 5 | 10 |
| not ROI | 121 | 869 | 990 |

For the illustrated subject (Figure 7), we observed |pSTS| = 10, |S_mod_ ∩ S_FC_| = 126, and an intersection of 5 parcels (Jaccard = 0.0382; precision = 0.0397; recall = 0.50). The enrichment was statistically significant by all tests (Fisher’s exact two-sided p = 4.38×10^−3^; hypergeometric right-tail p = 4.38×10^−3^; permutation p = 4.40×10^−3^), indicating that the ROI–group overlap is unlikely to occur by chance.

### Region of intersection of the key parcels across different modality models

We further localized the most important parcels for naturalistic viewing by another method - finding the area of intersection of the most important parcels for each of the seven different modality models and the 250 parcels found by piecewise “broken-stick” analysis, which can be seen in Figure 6B.

## Discussion

This study explored the hypothesis that the absolute value of FC can serve as a reliable indicator for identifying task-relevant brain parcels during naturalistic perception. The hypothesis is supported as the parcels identified by FC strength overlapped with previously established multimodal integration hubs and regions known to process signals during naturalistic movie viewing. Such confirmation allows FC-based scores to be used as a principled method for localizing the most relevant parcels for naturalistic perception. Beyond replication, this approach reveals previously unrecognized regions involved in multimodal perception and suggests new functional roles for regions already linked to naturalistic perception.

Our analyses revealed that a compact subset of parcels is significantly more important for multimodal perception than the rest of the brain regions. These parcels were identified using the average FC-based strength measure combined with the “broken-stick” analysis. The primary objective was to test whether FC-based interconnectedness is a robust marker of parcel importance for naturalistic perception and to gain insight into the spatial distribution of parcels supporting multimodal stimulus processing using analysis.

The hypothesis was supported: ablation analyses and cross-modality parcel intersections consistently identified regions whose predictive relevance aligned with known multimodal hubs—including pSTS, LOC, STG, and MTG—and other regions engaged by naturalistic movie viewing. This convergence validates the efficiency of the FC-based method for localizing task-relevant parcels.

The results further show that parcel contributions are highly non-uniform. A pronounced change in the curvature of the ablation curve emerged around the top ∼200 parcels, indicating a functional hierarchy. Models trained exclusively on these top-ranked parcels outperformed whole-brain models by ∼16.5%, despite using only a small subset of the brain. This finding demonstrates that focusing on task-critical parcels enhances prediction accuracy.

Interestingly, not all non-core parcels were equally irrelevant; some contributed negligibly or even detracted from prediction performance. This suggests that certain parcels may contain noise or unrelated variance that can hinder modeling when included indiscriminately.

These findings have several implications. First, they argue against treating the whole brain as equally informative for naturalistic perception tasks. Instead, predictive models should incorporate parcel-relevance weighting or targeted masking strategies guided by FC. Second, the approach can be extended to higher-resolution atlases to refine the localization of core integration hubs. Finally, the distributed yet structured nature of the identified network warrants further investigation to understand how these interconnected parcels coordinate during complex, real-world perception.

## Conclusion

In this study, we demonstrated that a brain parcel’s role in naturalistic perception can be reliably inferred from its absolute FC strength. Leveraging this principle, we introduced a “broken-stick” analysis to define a core multimodal integration subnetwork (∼250 parcels), that:

1. Enhanced predictive performance of multimodal encoding models,
2. Overlapped with known audiovisual and multimodal hubs (e.g., pSTS, STG, MTG, LOC), and,
3. Revealed new distributed components relevant to naturalistic perception(see supplementary section).

This approach provides a generalizable framework for identifying task-critical parcels in other domains and suggests that parcel-level relevance information can be integrated into future predictive model architectures to reduce training costs and boost accuracy.

By highlighting the unequal and connectivity-based contributions of parcels, this work advances our understanding of the network architecture underlying naturalistic perception localising S_sup_ a core of parcels relevant to the process of naturalistic viewing using a novel method and offers a principled path for refining brain-encoding models.

This study provides a method for attaining information about parcel’s relevance of the task through the analysis of an FC ablation curve. Future research venues may use this approach to study FC ablation curves for other tasks and identify most relevant parcels for those tasks. The found regions may be compared to known main regions in that area of research, advancing the understanding of various cognitive processes.

## Acknowledgements

This research was conducted by Viacheslav Fokin under the mentorship of PhD Arefeh Sherafati. We would like to acknowledge Lumiere Education for facilitating this research and providing edits to the manuscript. Many thanks to the CCN community for providing insightful comments.

## Supplementals

The parcels that were in the intersection of the most important parcels, as designated by all seven models, with the number of parcels participating in different brain regions of several brain atlases. Background means unlabeled parcels. The 7 model intersection parcels by Harvard, Oxford Cortical Structural Atlas

**Area**

~~~
Lateral Occipital Cortex; superior division
Superior Temporal Gyrus; posterior division
Lateral Occipital Cortex; inferior division
Occipital Pole
Occipital Fusiform Gyrus
Superior Frontal Gyrus
Temporal Pole
Background
Frontal Pole
Precentral Gyrus
Angular Gyrus
Middle Temporal Gyrus; posterior division
Supramarginal Gyrus; posterior division
Middle Temporal Gyrus; temporooccipital part
Precuneous Cortex
Superior Temporal Gyrus; anterior division
Heschl’s Gyrus (includes H1 and H2)
Inferior Frontal Gyrus; pars triangularis
Lingual Gyrus
Planum Polare
Temporal Occipital Fusiform Cortex
Frontal Opercular Cortex
Frontal Orbital Cortex
Inferior Frontal Gyrus; pars opercularis
Inferior Temporal Gyrus; anterior division
Juxtapositional Lobule Cortex (formerly Supplementary Motor Cortex)
Middle Frontal Gyrus Planum Temporale
The 7 model intersection parcels by Destrieux Cortical Atlas
~~~

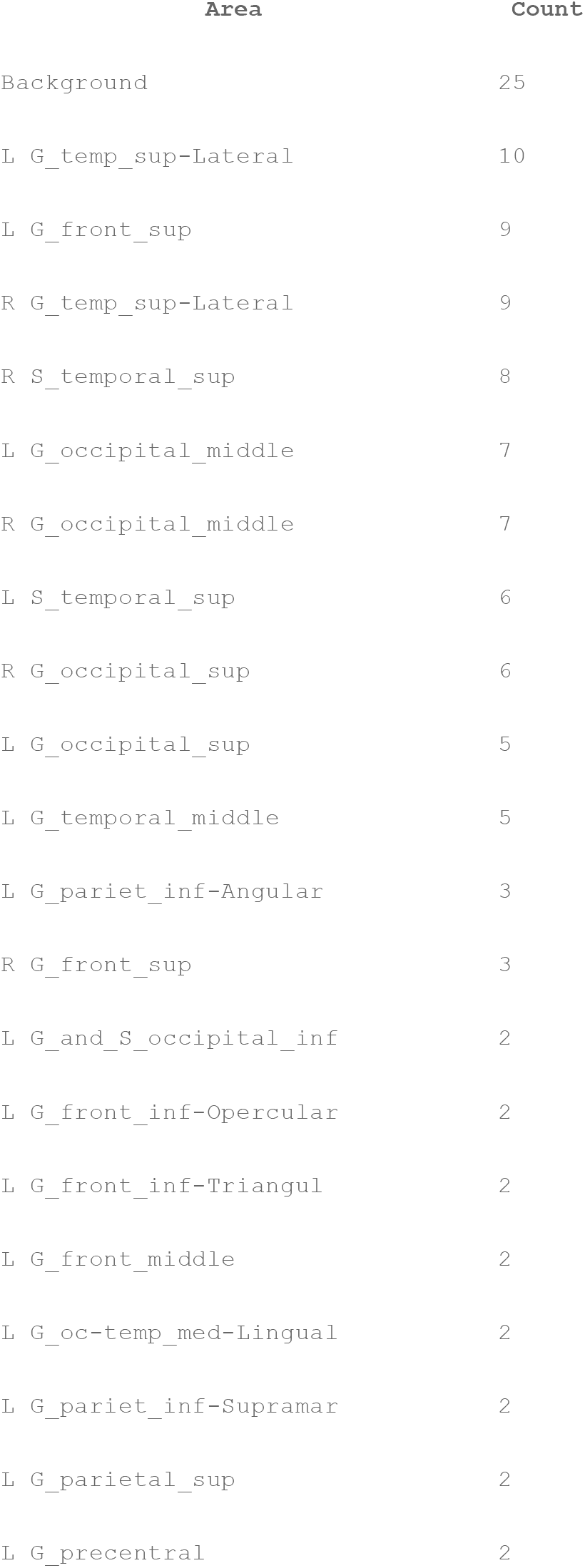
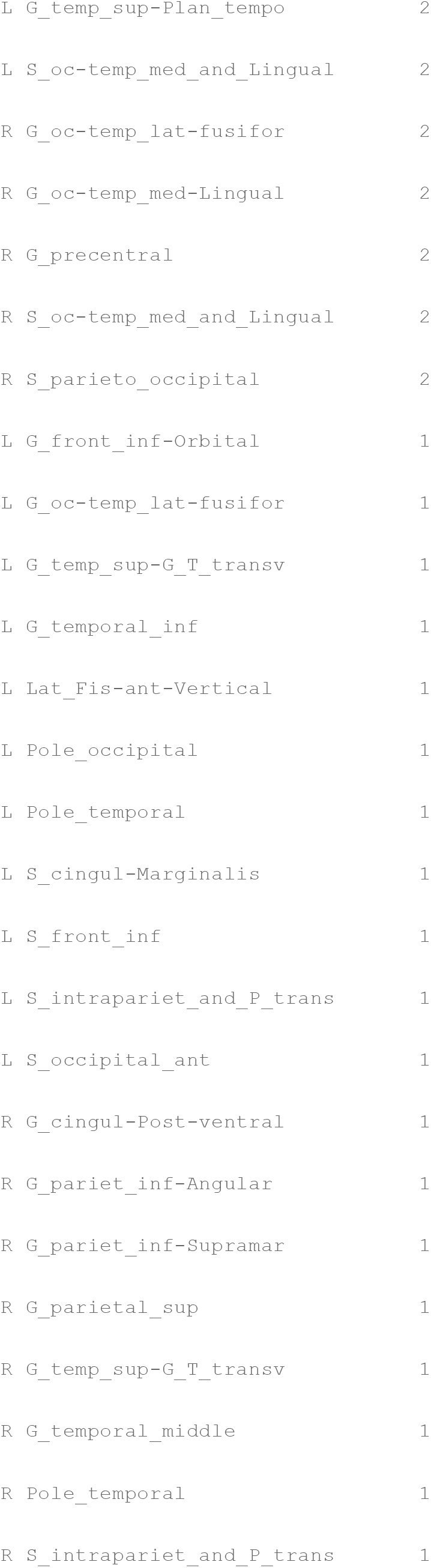

The 7 model intersection parcels by Destrieux Cortical Atlas
| Area | Count |
| --- | --- |
| Background | 25 |
| L G_temp_sup-Lateral | 10 |
| L G_front_sup | 9 |
| R G_temp_sup-Lateral | 9 |
| R S_temporal_sup | 8 |
| L G_occipital_middle | 7 |
| R G_occipital_middle | 7 |
| L S_temporal_sup | 6 |
| R G_occipital_sup | 6 |
| L G_occipital_sup | 5 |
| L G_temporal_middle | 5 |
| L G_pariet_inf-Angular | 3 |
| R G_front_sup | 3 |
| L G_and_S_occipital_inf | 2 |
| L G_front_inf-Opercular | 2 |
| L G_front_inf-Triangul | 2 |
| L G_front_middle | 2 |
| L G_oc-temp_med-Lingual | 2 |
| L G_pariet_inf-Supramar | 2 |
| L G_parietal_sup | 2 |
| L G_precentral | 2 |
| L G_temp_sup-Plan_tempo | 2 |
| L S_oc-temp_med_and_Lingual | 2 |
| R G_oc-temp_lat-fusifor | 2 |
| R G_oc-temp_med-Lingual | 2 |
| R G_precentral | 2 |
| R S_oc-temp_med_and_Lingual | 2 |
| R S_parieto_occipital | 2 |
| L G_front_inf-Orbital | 1 |
| L G_oc-temp_lat-fusifor | 1 |
| L G_temp_sup-G_T_transv | 1 |
| L G_temporal_inf | 1 |
| L Lat_Fis-ant-Vertical | 1 |
| L Pole_occipital | 1 |
| L Pole_temporal | 1 |
| L S_cingul-Marginalis | 1 |
| L S_front_inf | 1 |
| L S_intrapariet_and_P_trans | 1 |
| L S_occipital_ant | 1 |
| R G_cingul-Post-ventral | 1 |
| R G_pariet_inf-Angular | 1 |
| R G_pariet_inf-Supramar | 1 |
| R G_parietal_sup | 1 |
| R G_temp_sup-G_T_transv | 1 |
| R G_temporal_middle | 1 |
| R Pole_temporal | 1 |
| R S_intrapariet_and_P_trans | 1 |

